# Eye movements organize excitability state, information coding and network connectivity in the human hippocampus

**DOI:** 10.1101/2024.12.12.628236

**Authors:** Marcin Leszczynski, Elizabeth Espinal, Elliot Smith, Catherine Schevon, Sameer Sheth, Charles E. Schroeder

**Author notes:** Correspondence to: Dr. Marcin Leszczynski or Dr. C.E. Schroeder Department of Psychiatry, Columbia University College of Physicians and Surgeons, 1051 Riverside Drive Kolb Annex Rm 561, New York, NY 10032, <u>Email:</u>.

## Abstract

Natural vision is an active sensing process that entails frequent eye movements to sample the environment. Nonetheless vision is often studied using passive viewing with eye position held constant. Using closed-loop eye-tracking, with saccade-contingent stimulation and simultaneous intracranial recordings in surgical epilepsy patients, we tested the critical role of eye movement signals during natural visual processing in the hippocampus and hippocampal-amygdala circuit. Prior work shows that saccades elicit phase reset of ongoing neural excitability fluctuations across a broad array of cortical and subcortical areas. Here we show that saccade-related reset systematically modulates neuronal ensemble responses to visual input, enables phase-coding of information across the saccade-fixation cycle and modulates network connectivity between hippocampus and amygdala. The saccade-fixation cycle thus emerges as a fundamental sampling unit, organizing a range of neural operations including input representation, network connectivity and information coding.

**One-Sentence Summary:** Saccade-fixation cycle: a fundamental sampling unit, organizing input representation, information coding and network activation

## Introduction

In a natural environment, sensing is an active process. Most sensory inputs that enter brain are acquired actively by motor sampling routines (*1–5*). In visual sensing, for example, we actively seek visual information using saccadic eye movements and head/body rotations to scan the environment and seek out objects of interest. The process of active sensing is conserved across sense modalities and species, encompassing whisking (*1*, *3*) and sniffing in rodents (*6*, *7*), as well as tactile (*8*, *9*) and saccadic sampling in primates (*4*, *5*, *10–12*).

Critical to *visual* active sensing is the fact that a motor act, the saccade, causes stimulation of sensory receptors, initiating the inflow of activation that propagates through sensory systems. Earlier studies of the effects of saccades in the dark (*13–21*), and more recent studies that minimize saccade-related visual input by various means (*22–25*) confirm the proposition that non-retinal “corollary discharge” signals, generated in parallel to saccades, modulate the excitability of neurons throughout the visual pathways.

Intriguingly, saccadic modulation spreads out into classic auditory and somatosensory areas (*26*) as well as supramodal areas like the hippocampus (*24*, *25*, *27–31*) and the thalamus (*32*). Network connectivity is modulated across the saccade-fixation cycle with increased top-down interactions from the frontal eye fields to sensory areas during saccades and reversed, bottom-up interaction after fixation onset (*26*). These findings align with an earlier proposition advanced at the level of interlaminar cell circuitry in V1 (*23*).

How do saccadic signals modulate neural activity? Several lines of evidence (*5*, *19*, *23*, *33*, *34*) suggest that nonretinal signals, corollary to saccade generation, enhance sensory processing. We and others proposed that sacades enhanced sensory processing by resetting the phase of ongoing excitability fluctuations (aka oscillations; (*4*, *5*, *24*, *25*, *29*)). According to this proposition, the phase reset of an oscillation, with period corresponding to the duration of a typical 200-300 ms saccade-fixation cycle, aligns the high-excitability phase of that oscillation with the arrival of visual input in primary visual cortex (V1) (*19*, *23*), thus amplifying the neuronal response to visual input. Similar phase reset occurring across levels of the visual hierarchy could facilitate passage of information up the system. In support of this idea previous studies observed an increase in phase coherence following saccade across the visual hierarchy all the way up to the hippocampus (*24*, *25*, *29*) and directly connected thalamic nuclei (*32*).

The goal of this study was to further explore the role of saccade-related signals in sensory processing. First, we hypothesized that the saccade-related signals reset of ongoing excitability fluctuations (oscillations), reorganizes neuronal ensembles to a high excitability state, which should enhance the input evoked response. To test this prediction, we compared evoked responses elicited by the same stimuli presented immediately after saccades (active condition) with stimulus evoked responses during sustained fixation (passive condition). Second, the visual active sensing model predicts that neuronal excitability oscillates across the saccade-fixation cycle. We tested this prediction by examining the magnitude of stimulus evoked responses as a function of the lag between stimulus onset and saccade onset. Third, recent computational work (*35–37*) led us to predict that the low excitability phase occurring later in the saccade-fixation cycle may enhance selectivity of information coding, perhaps by depressing noise correlations in local neuronal populations. We thus examined the specificity of stimulus feature encoding across the saccade-fixation cycle. Finally, the visual active sensing model predicts that saccade-related neuronal excitability modulation enhances the flow of information through brain circuitry. We tested this idea by comparing the strength of hippocampal-amygdala connectivity during passive versus active visual conditions.

The analysis focused on hippocampus as it is a supramodal structure that exhibits both oscillatory phase modulation (*24*, *25*, *28–31*, *38–40*) and phase-coding of information (*39*) during active sensing. We also monitored the hippocampal-amygdala circuit to index saccadic effects on network interactions. Building on the prior active vision work, we developed a closed-loop eye tracking/stimulus-presentation routine in combination with intracranial EEG recordings from the hippocampus and amygdala of surgical epilepsy patients to investigate how the physiology of visual processing differs between active and passive viewing of the same visual stimulus. By systematically exploring neural activity across the saccade-fixation cycle during active vision, we aimed to: 1) evaluate the degree to which **passive** visual processing approximates or differs from visual **active** sensing and 2) define consequences of saccadic phase reset for processing of sensory inputs.

Our findings support a model in which active vision differs fundamentally from passive vision in three key physiological dimensions. First, saccadic phase reset enhances hippocampal neuronal response to inputs entering the system following fixation. The dynamics of neural excitability is complementary to that known from primates’ V1. Second, the systematic excitability oscillation tied to the saccade-fixation cycle provides a potential substrate for phase coding of information. Finally, active saccadic sampling increases the gain of network interactions between hippocampus and amygdala. Our findings support a model in which the saccade-fixation cycle is a fundamental unit of information processing, one that is excluded by studying the system in a passive vision paradigm with eye position held constant or uncontrolled.

## Results

### Active vision enhances neural response in the human hippocampus

Participants performed a modified delayed match-to-sample task, where each trial commenced with a one-second presentation of an image centrally (in the active condition) or within one of four designated locations (in the passive condition; Fig. 1A-B), requiring memorization. Following image offset, participants performed either an active visual search involving saccadic eye movements across the screen (in the active condition) or a passive visual search which require maintaining fixation (in the passive condition). Note that image was displayed (presentation duration of 17 ms) only if sustained fixation was detected (passive) or if participants changed their gaze location to a new hot-spot (active). At randomly selected points (from a range of 4-14 images) the search was stopped and participants were prompted to determine whether the last image matched the initial one (balanced and randomized responses). We analyzed data from the visual search part. It is important to note that the task was designed to ensure comparable levels of memory and attentional demand across both conditions, with the primary distinction lying in the necessity to either relocate gaze (active condition) or sustain fixation (passive condition). Overall accuracy in the delayed-match-to-sample task was high, reaching 81.43% in the active condition and 81.07% in the passive condition (Wilcoxon sign-rank test; p = 0.73; z = 0.33; N = 7). Response times were long in both conditions (speed was not encouraged) but were shorter during active compared to passive vision (2.41 sec vs. 4.37 sec; Wilcoxon sign-rank test; p = 0.01; z = 2.36; N = 7; see Fig. 1D). Additionally, the inter-stimulus interval remained consistent across both conditions, with frequencies of 1.38 Hz in the active and 1.28 Hz in the passive vision (Wilcoxon sign-rank test; p = 0.13; z = 1.52; N = 7).

**Figure 1:**
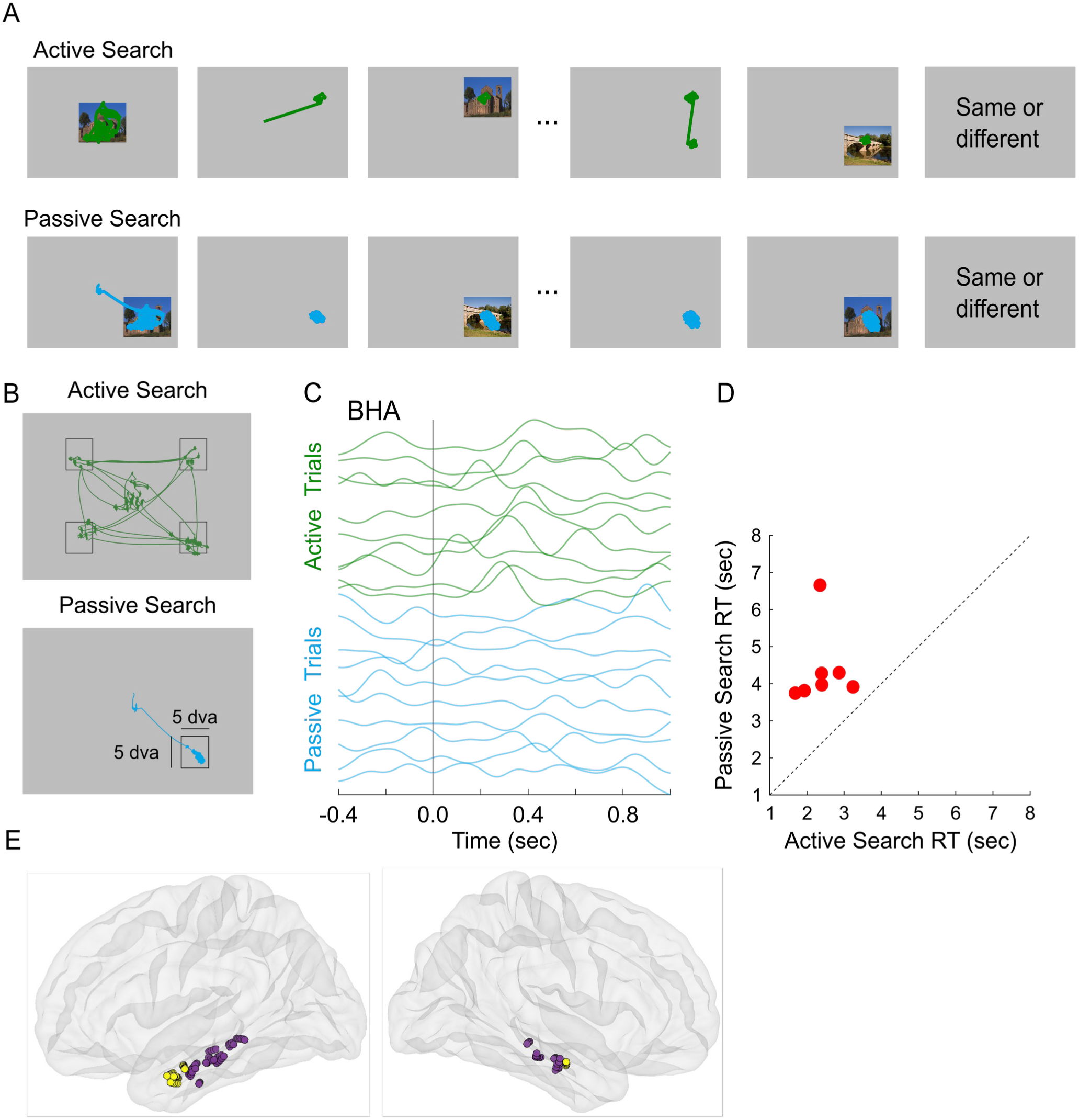
Task, behavior and electrode locations. **(A)** Trial sequence in the active (upper row) and passive (lower row) search conditions, with eye-tracking traces illustrating oculomotor behavior. In the active condition, participants fixated on a central image (5 degrees of visual angle) for 1 second, then searched for images at one of four pre-designated locations. Upon fixating on a location (e.g., upper right corner), an image was presented for 17 ms. Participants continued to the next location until prompted to identify if the last image matched the initial one. In the passive condition, participants fixated on an image in one of the quadrants for 1 second. After the image disappeared, they kept their eyes on this location while a sequence of images was presented for 17 ms each. The inter-stimulus interval and stimuli were matched to the active condition. This experiment was also designed to equate attention effects across both conditions, using the same set of images, comparable inter-stimulus-interval and comparable delayed match to sample task. Participants were prompted to identify if the last image matched the initial one. **(B)** Example eye trace from active (upper panel) and passive (lower panel) visual search sequences (see panel A) with 5 dva ‘hot-spot’ that superimposed for illustration. Note that squares in the figure are illustrative only and were not shown to participants **(C)** Ten examples of single-trial BHA from a hippocampal channel during active (green) and passive (blue) search. X-axis: time relative to stimulus onset. Stimulus “onset” and “offset” refer to turning the stimulus on and off, respectively (i.e., the beginning and the end of stimulus presentation). **(D)** Behavioral performance in active (x-axis) and passive (y-axis) search. For each participant we plotted response time to the question “Was the last image was old or new?” Note all patients showed shorter response times in the active compared to passive visual search. Accuracy was high and did not differ across conditions. **(E)** Anatomical distribution of hippocampal (n = 76 channels across 7 patients; violet circles) and amygdala channels (n = 36; across 4 patients; yellow circles) which were transformed onto an average brain. Left and right panels: lateral view (left and right, respectively).

To understand physiology of active vision we studied field potentials elicited by stimuli during active and passive vision. We analyzed data from the human hippocampus because previous studies identified the medial temporal lobe and hippocampus as part of a brain-wide network modulated during active vision (*24*, *25*, *28–32*).

We probed local neural processes using Broadband High-frequency Activity (BHA; 70-150 Hz) which reflects a mixture of fast dendritic potentials and action potentials and closely correlates with multi-unit activity (*41*, *42*). We also studied event-related potentials (ERPs) as well as inter-trial phase coherence (ITC) and power (1-30 Hz). We used these to measure the influence of saccades on the neural response elicited by visual stimuli. Comparing neural activity in two conditions with the same visual input presented either following a saccade or sustained fixation (i.e., active and passive search, respectively), holds visual stimulation constant, allowing us to isolate effects due of saccadic modulation of local neuronal activity.

Figure 1C displays ten examples of single-trial stimulus onset related BHA recorded from the human hippocampus during active and passive vision. We noted that the magnitude of neural response is higher in active condition trials. Directly comparing hippocampal BHA (n = 76 channels, 7 patients), confirmed the above impression of enhanced response in active condition (Fig. 2A; unless stated differently we used Wilcoxon sign-rank test controlled for multiple comparisons with Benjamini-Hochberg procedure; all p < 0.05; all z > 1.96; n = 76). The biggest difference was 300-400 ms after stimulus presentation, which corresponds to the strongest sensory elicited hippocampal response (*43*, *44*). Similarly, ERPs showed faster onset and higher magnitude in active vision (Wilcoxon sign-rank test; all p < 0.05; all z > 1.96 controlled for multiple comparisons with Benjamini-Hochberg procedure; n = 76). We also noted that the ERP in active condition transiently decreased in a later time window around 600 ms after stimulus onset (Fig. 2B; Wilcoxon sign-rank; all p < 0.05; all z > 1.96 controlled for multiple comparisons with Benjamini-Hochberg procedure; n = 76).

**Figure 2:**
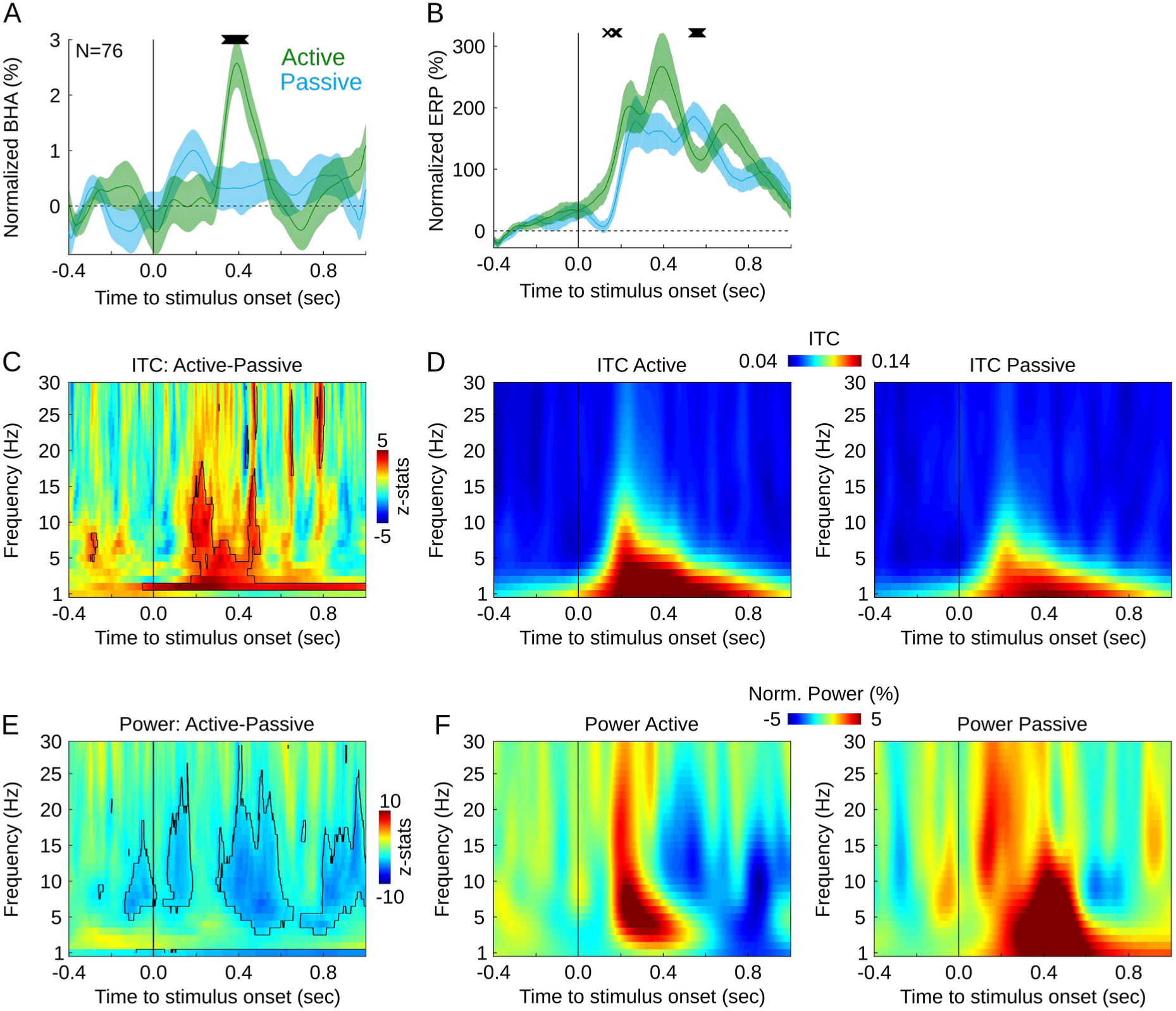
Neural activity during active and passive vision. Line plots show BHA **(A)** and ERP **(B)** measures in the hippocampus elicited by the same set of images during active (green) and passive (blue) vision. Y-axis: magnitude of BHA (A) and ERP (B) relative to the prestimulus baseline. Shading reflects standard error of the mean (SEM). Black markers over the plot indicate time points with significant difference between active and passive. Color maps show results of ITC **(C)** and power **(E)** comparison between active and passive conditions (z-statistics from Wilcoxon sign rank test). Warm color indicates higher values in active. Contours in D, F: time-frequency points of significant difference between conditions. **(D, F)** present ITC (D) and power (F) in active (left panels) and passive (right panels) condition. Power, as with BHA and ERPs is normalized relative to the prestimulus baseline. Y-axes: frequency (panels D-G). All x-axes: time relative to stimulus onset except panel C which shows frequency. All horizontal lines: stimulus onset. All results FDR controlled for multiple comparisons with the Benjamini-Hochberg procedure.

Because an ERP consists of a mixture of the phase reset and evoked component (*45–48*) (i.e., power increase) these observed ERP modulations may originate from either of these two elements. To further explore how saccades influence sensory evoked potential, we examined stimulus-related phase coherence and power in active and passive condition. We found increased phase coherence associated with power decrease in the active condition (Wilcoxon sign-rank; all p < 0.05; all z > 1.96 controlled for multiple comparisons with Benjamini-Hochberg procedure; n = 76). Specifically, we observed two ITC components which, for descriptive purposes, we refer to as: 1) “early” – peaking around 200 ms after stimulus onset, centered at alpha frequency range (5-15 Hz); 2) “late” – peaking around 300-400 ms after stimulus, centered at theta frequency range (2-5 Hz; see Fig. 2C-D for exact statistics results). We also observed power modulations during active condition detectable in a number of frequencies across theta, alpha and beta ranges and oscillating at theta/alpha frequencies (Fig. 2E-F; Wilcoxon sign-rank; all p < 0.05; all z > 1.96 controlled for multiple comparisons with Benjamini-Hochberg procedure; n = 76). As observed with BHA, these effects peaked about 200 – 450 ms after the stimulus onset. Control analyses confirm that all the above effects are unchanged when we account for potential clustering among our measurements with linear mixed effects models using experimental condition as a fixed effect and treating participant as a random effect (see Supplemental Information).

These findings indicate that sensory stimuli during active vision elicit stronger and faster neuronal responses as indexed by BHA and ERP. Note, that a combination of increased ITC and decreased power show that the difference between active and passive vision is qualitative: active vision is not only a stronger version of passive vision but entails fundamentally different neural processes.

Importantly, the current results are unlikely to be explained by unequal allocation of attentional or memory resources (see Discussion). Overall, these data support a model in which a signal generated in parallel to saccades resets neural oscillations to a higher excitability state around the time of local response to sensory input. Passive vision, by eliminating saccadic eye movements (and the associated phase reset), differs fundamentally from active vision.

### Dynamics of excitability across the saccade-fixation cycle

These results support the model in which saccade-related signals modulate the neural response to visual input (*19*, *23*, *26*, *32*). We further hypothesized that these saccade-related signals initiate cyclic changes in neural excitability following fixation (*5*). To test this, we systematically probed the system’s excitability across the saccade-fixation cycle by presenting stimuli at 50, 100, 150, 200 ms time lag relative to fixation onset. We measured BHA at these four distinct phases and found that the magnitude of stimulus-evoked BHA varied across the saccade-fixation cycle, rather than remaining constant (Fig. 3). We identified three time windows in which BHA differed across four conditions (Kruskal-Wallis test, all p < 0.01 controlled for multiple comparisons using Benjamini-Hochberg correction; see Methods). The biggest modulation of the BHA was in an early time window (170 – 250 ms after stimulus onset). Specifically, BHA magnitude increased from lag 1 to lag 2 (z = 3.46, p < 0.001; Wilcoxon sign rank test; n = 76), decreased from lag 2 to 3 (z = 2.37, p = 0.01; Wilcoxon sign rank test; n = 76) and increased again from lag 3 to 4 (z = 3.47, p < 0.001; Wilcoxon sign rank test; n = 76). When comparing early BHA in passive vision to that observed at different lags during active vision, we observed that BHA in lags 1 and 3 was reduced compared to passive vision (both z > 3.51, both p < 0.001; Wilcoxon sign rank test; n = 76), while BHA in lags 2 and 4 did not differ from that in passive vision (both z < 1.64, both p > 0.09; Wilcoxon sign rank test; n = 76). This shows that hippocampal BHA is initially suppressed after fixation onset, followed by oscillatory patterns of enhancement and suppression with a period of approximately 100 ms (Fig. 3B).

**Figure 3:**
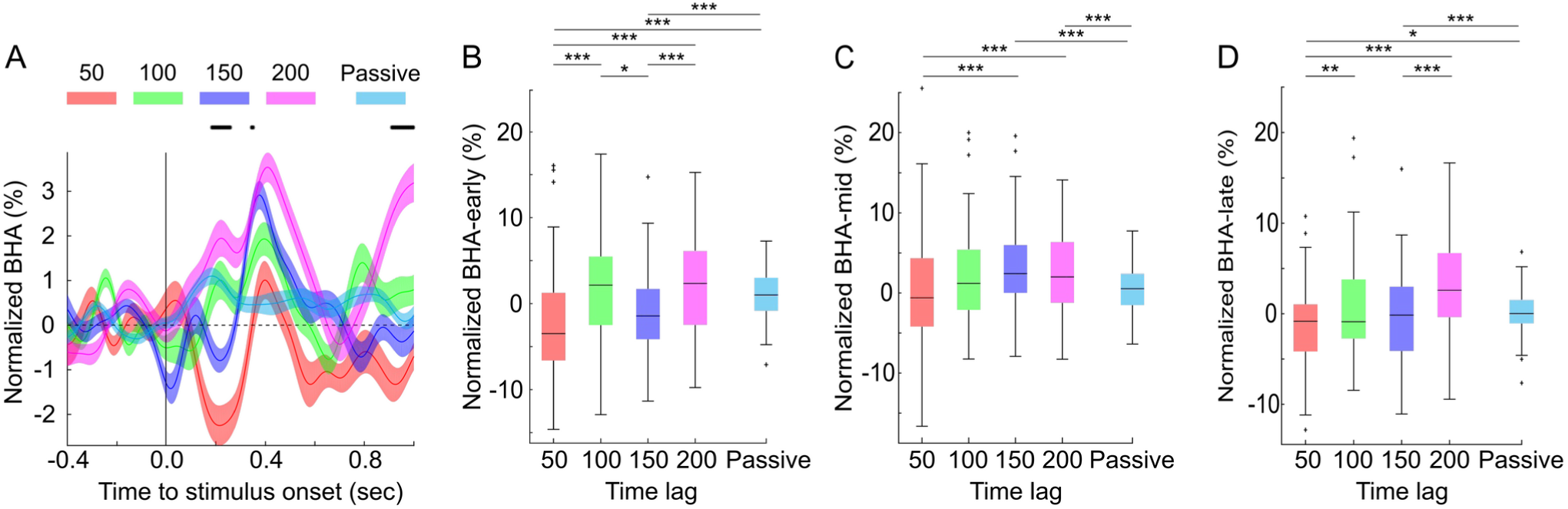
Neural excitability in the human hippocampus across the saccade-fixation cycle. **(A)** BHA elicited during four active vision conditions (i.e., for stimuli presented 50, 100, 150, 200 ms following fixation onset; red, green, dark blue, magenta respectively) and passive vision (light blue). Shading reflects SEM. X-axis show time, Y-axis show magnitude of BHA relative to pre-stimulus baseline. The vertical line indicates stimulus onset. Small crosses over the plot indicate time points when BHA differed across conditions (Kruskal-Wallis test, all p < 0.01 controlled for multiple comparisons using Benjamini-Hochberg correction). **(B, C, D)** Box plots show average BHA during the early (170 – 250 ms) and middle (340 – 350 ms) and late (900-1000 ms) time window, respectively. Box plots indicate 25^th^, median and 75^th^ percentile, whiskers extend to extreme values not considered outliers while outliers are marked with crosses. Star symbols show p-values from a pairwise sign rank test: *,**,*** reflect p < 0.05, 0.01 and 0.005, respectively.

We also observed two later windows where BHA magnitude deviated from uniformity across the saccade-fixation cycle (Fig. 3 A-D; 340 – 350 ms and 900-1000 ms after stimulus onset; both identified with Kruskal-Wallis test, all p < 0.01 controlled for multiple comparisons using Benjamini-Hochberg correction; see Methods). In the 340 – 350 ms window, BHA magnitude steadily increased across time lags (Fig. 3C) with long-lag responses greater than those found in passive vision (below we report Wilcoxon sign rank test results; all n = 76): BHA increased from lag 1 to lag 3 (z = 3.45, p < 0.001) and lag 4 (z = 2.96, p = 0.003) but not from lag 1 to lag 2 (z = 1.72, p = 0.083). Additionally, BHA at the longest lags (lag 3: z = 4.45, p < 0.001; lag 4: z = 2.89, p = 0.003) was greater than that observed during passive vision, whereas shortest lags showed no significant difference (lag 1: z = 0.76, p = 0.44; lag 2: z = 1.88, p = 0.058). This indicates that visual stimuli elicit a stronger BHA component as time from fixation onset increases.

In the late BHA window (900-1000 ms), we observed a modulation pattern similar that in the early window: BHA magnitude increased from lag 1 to lag 2 (z = 2.56, p = 0.01; Wilcoxon sign rank test; n = 76) and from lag 3 to lag 4 (z = 3.70, p < 0.001; Wilcoxon sign rank test; n = 76), with no significant difference between lag 2 and lag 3 (z = 0.78, p = 0.43; Wilcoxon sign rank test; n = 76). Comparing BHA in passive vision with that observed at different lags during active vision, we found that BHA in lag 1 decreased relative to passive vision (z = 2.21, both p = 0.02; Wilcoxon sign rank test; n = 76), while BHA in lag 4 increased (z = 4.24, both p < 0.001; Wilcoxon sign rank test; n = 76). BHA magnitude did not differ significantly from passive vision in lags 2 and 3 (both z < 0.92, both p > 0.35; Wilcoxon sign-rank test; n = 76).

Altogether, these results show that the magnitude of visual evoked potential in the hippocampus is suppressed right after fixation and then oscillates between enhancement and suppression with a period of about 100 ms. Additionally, the magnitude of visual evoked potential in the hippocampus steadily increases following fixation onset. This shows that hippocampal neural excitability fluctuates at two different rates across the saccade-fixation cycle (Fig. 3B-C): a faster cycle of approximately 100 ms (Fig 3B) embedded into a slower steady increase (Fig. 3 C,D). These dynamic patterns are missing in passive viewing conditions.

### Coding of information across the saccade-fixation cycle

We considered two hypothetical ways that modulation of BHA response amplitude could regulate the encoding of visual information across the saccade-fixation cycle. Information encoded during a high excitability phase of the saccade-fixation cycle (indexed by a stronger BHA response) may have greater salience in ongoing processing, due to an amplified neuronal response. Alternatively, the low excitability phase (indexed by weaker BHA response) may enhance selectivity of information coding, perhaps by depressing noise correlations in local neuronal populations (*35–37*). Additionally, hippocampus receives convergence of dorsal and ventral pathway input (*49*) which carry preferential representation of thalamic magnocellular (M) and parvocellular (P) input, respectively (*50*). As, M inputs travel into the system more rapidly than P inputs (*50*), we hypothesized that M-mediated features like intensity and gross form/edge would impact the hippocampal visual representation at an earlier phase of the fixation-saccade cycle than P-mediated features like color and orientation. In contrast, previous psychophysics studies found that coarser elements (i.e., intensity) are represented closer to fixation onset, followed by finer elements (i.e., edges) (*51–54*). The null hypothesis is that the amount of information is uniformly distributed across the saccade-fixation cycle.

To investigate these possibilities, we conducted two sets of analyses. We utilized representational similarity analysis (RSA) to assess the quantity of visual information present in field potentials and a decoding approach (Supplementary Text). Both of these analyses show that information is differentially encoded across the saccade-fixation cycle. Specifically, using RSA we probed the extent to which evoked patterns track visual features (i.e., intensity, edges, saliency, orientation and color) in the stimulus space across the saccade-fixation cycle (see Fig. 4A and Methods for the analysis overview). If field potentials contained more information about visual features at any particular phase of the saccade-fixation cycle, we expected the RSA to be non-uniform across the cycle. We also expected that different information (i.e., features) may be preferentially encoded at different phases of the cycle.

**Figure 4:**
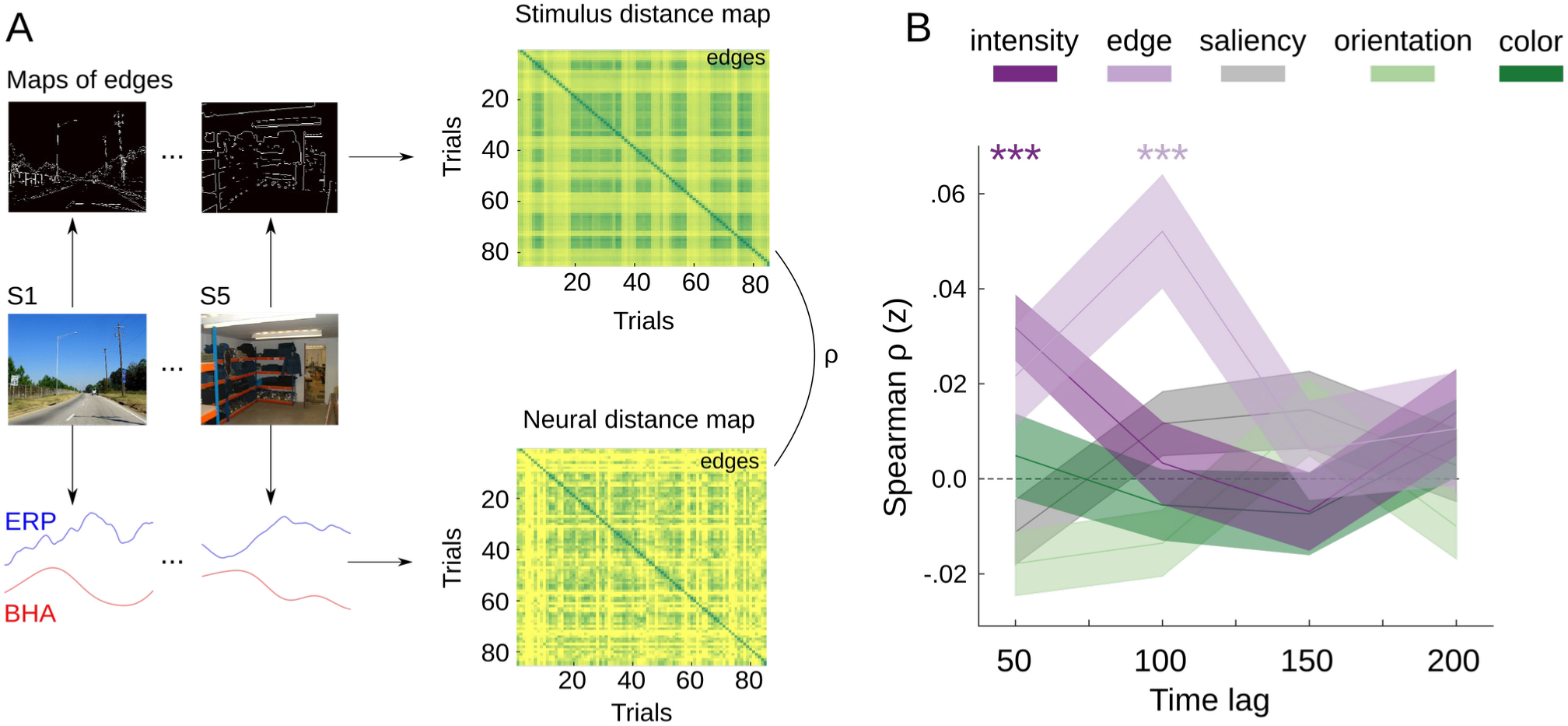
Coding of information across the saccade-fixation cycle. **(A)** We investigated the ability of neural activity to track stimulus features (edges, saliency, orientation, intensity, and color) throughout the saccade-fixation cycle. Using Euclidean distance matrices, we quantified the similarity between pairs of stimuli and pairs of evoked potentials at each time lag. The neural signals consist of 200 ms long snippets of BHA and ERP centered at the time of strongest sensory evoked potential (i.e., 300-500 ms after stimulus onset). By correlating the stimulus distance matrix with the neural distance matrix, we assessed the extent to which neural patterns preserved stimulus-specific information. Increased correlation coefficients indicated higher amount of information about the visual feature (i.e., better feature tracking by the neural data). **(B)** Spearman’s rho (Fisher’s z transformed) between stimulus and neural distance maps was computed for each feature and time lag. Y-axis: Spearman’s rho. X-axis: four time lags. Shading reflects SEM and star symbols indicate pairwise sign rank test; p-values: *** denote p_corrected_ < 0.005.

Our findings support the phase coding hypothesis, showing that the degree to which neural data tracked stimulus features varied across the saccade-fixation cycle. We found that the RSA differed across four time lags for the edges, intensity and salience (all H(3) > 10.27, all p < 0.05; Kruskal-Wallis test; n = 76), but not for the color and orientation (all H(3) < 7.01, p > 0.07; Kruskal-Wallis test; n = 76). To account for the potential influence of trial numbers on correlation coefficients, we conducted a control analysis where we recalculated RSA after matching the number of trials across conditions. Our control analysis confirmed the initial results: again, neural tracking of stimulus features varied across lags for edges, intensity and overall salience (all H(3) > 8.02, all p < 0.05; Kruskal-Wallis test; n = 76), but not for the color and orientation (all H(3) < 7.40, p > 0.06; Kruskal-Wallis test; n = 76). These results demonstrate that at least some types of sensory information are differentially encoded across time during the saccade-fixation cycle.

To investigate the preferred phases associated with specific features, we performed a series of one-tailed Wilcoxon sign-rank tests, with careful control for multiple comparisons using Benjamini-Hochberg correction. For each feature and lag, our null hypothesis stated that the RSA at a given lag is not higher than zero (for similar approach see (*55–57*)). Our analysis showed that information about intensity was selectively represented at the earliest lag (i.e., 50 ms; p < 0.005; z = 4.21; Wilcoxon sign rank test; n = 76), followed by information about edges selectively represented at 100 ms (p < 0.005; z = 3.74; Wilcoxon sign rank test; n = 76; see Fig. 4B). No other features reached significant correlation at any lag. Note that we used a-priori the one-tailed test but a two-tailed version of the analyses reproduced our main results. To ensure that differences in image features across lags did not account for these results, we explicitly compared feature distributions across conditions and found no differences (all p > 0.13; all chi-sq < 5.55; Kruskal-Wallis test; N = 7). These results suggest that information about intensity and edges is preferentially represented at specific phases of the saccade-fixation cycle. Importantly, information about intensity is represented closer to the saccade and is followed by information about edges supporting the idea of sequential coarse-to-fine analysis of visual input across the saccade-fixation cycle (*51–54*).

### The hippocampus disconnects from amygdala to support sensory encoding in active vision

We also asked whether active vision modulates patterns of inter-regional connectivity in the hippocampal-amygdala circuit, which is known for its extensive reciprocal connections (*58*). Prior studies have demonstrated changes in phase synchronization in this circuitry during exposure to emotionally salient stimuli (*59*), with theta synchronization and alpha desynchronization facilitating mnemonic discrimination (*60*) likely through bias of attention to emotionally laden features during encoding. Because active vision enhances encoding of sensory stimuli in the absence of emotional saliency, we explored the possibility that theta synchronization and alpha desynchronization pattern also supports encoding of other than emotional forms of saliency (e.g., images acquired by voluntary gaze allocation).

First, we explored local neural activity in the amygdala (36 channels across 5 patients) during active and passive vision. Our analyses revealed increased ITC that peaked around 400 ms post-fixation and centered at 4 Hz (Fig. 5A). Interestingly, the overall spectro-temporal profile of the ITC in the amygdala resembled that observed in the hippocampus. Unlike the hippocampus, we detected no difference in ITC between active and passive vision (Wilcoxon sign-rank test; all p > 0.05; all z < 1.96; n = 36). Although BHA appeared decreased in active condition around the stimulus onset this effect did not survive correction for multiple comparisons, likely due to small sample size (Fig. 5B; Wilcoxon sign-rank test; all p > 0.05; all z < 1.96; n = 36). We also observed no difference in low frequency power (<30 Hz) or ERP amplitude (Fig. 5C).

**Figure 5:**
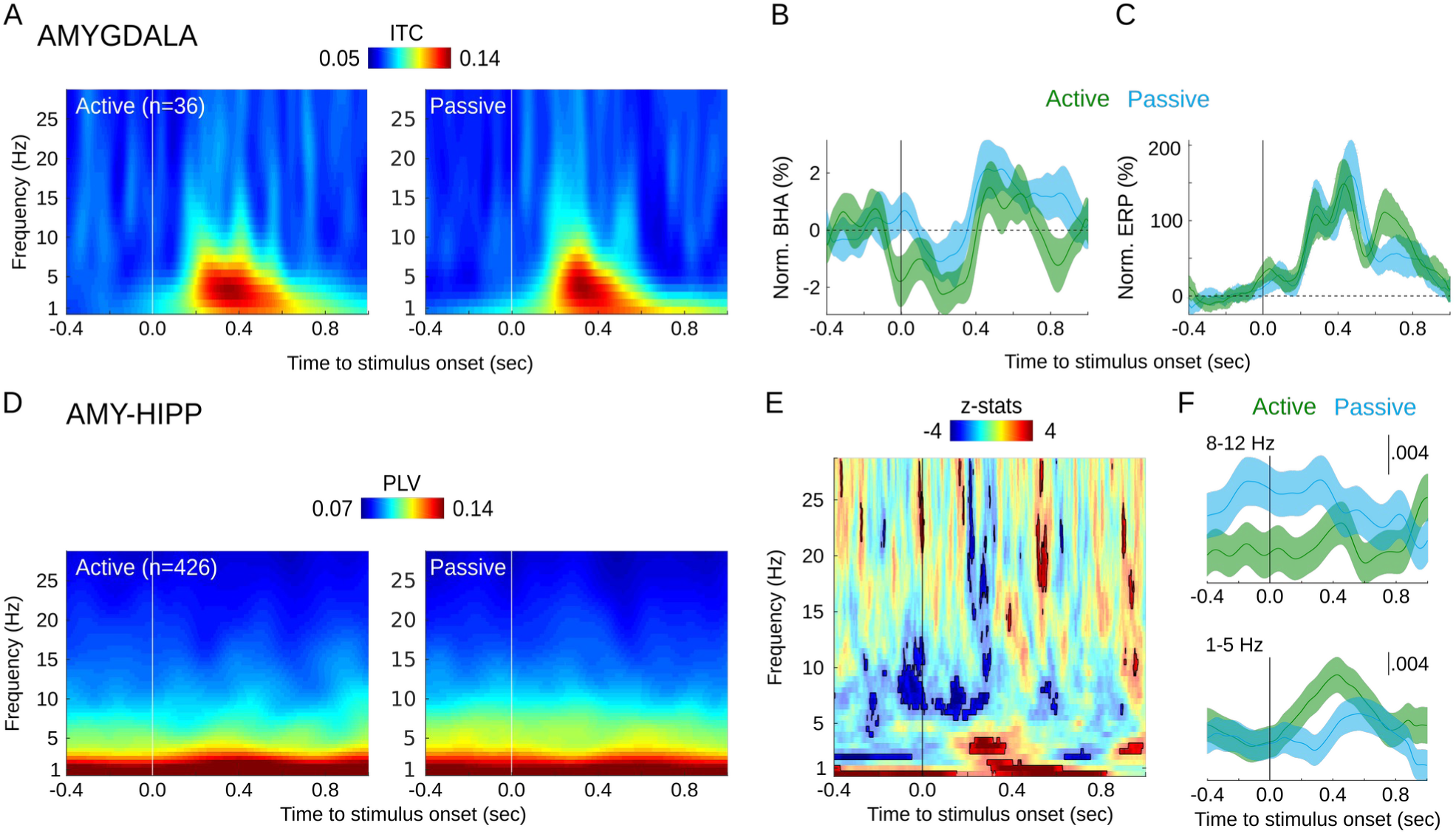
Network connectivity during active and passive vision. **(A)** Color maps show averaged stimulus-locked ITC in amygdala during active (left) and passive vision (right panel; n = 36 channels; 5 patients; x-axis: time, y-axis: frequency). Normalized BHA **(B)** and ERP **(C)** time locked to stimulus onset. Shading reflects SEM. **(D)** Color maps show averaged stimulus onset locked PLV between amygdala and hippocampus during active (left) and passive vision (right panel; n = 426 pairs of channels). **(E)** Z-statistics from a sign rank test comparing PLV between active and passive conditions. Negative values (blue) indicate time-frequency points where PLV is stronger during passive vision; positive values (red) indicate the reverse (i.e., stronger PLV during active vision). Contours depict significant time-frequency points (p < 0.05 controlled for multiple comparisons with the Benjamini-Hochberg procedure). **(F)** Averaged PLV in alpha (8-12 Hz; upper panel) and theta range (1-5 Hz; lower panel) where we found significant differences between active and passive vision. Vertical lines in all plots mark stimulus onset.

To investigate the impact of active vision on inter-regional network interactions, we employed the phase locking value (PLV) (*61*), a measure of phase consistency between two signals. Analyzing paired recordings from the hippocampus and amygdala we observed that the PLV decreased during active vision in the lower alpha range (i.e., 8-12 Hz) prior to and after fixation onset – here a proxy for image onset (Fig. 5D-E; Wilcoxon sign-rank; all p < 0.05; all z > 1.96 controlled for multiple comparisons with Benjamini-Hochberg procedure; n = 426). Importantly, this effect begun before fixation onset (and even before saccade execution) and continued after fixation onset. Control analyses showed that this dynamic did not change with the time lag. Additionally, we observed that the PLV increased during active condition in lower frequencies (1-5 Hz) around 300-500 ms after stimulus onset (Wilcoxon sign-rank; all p < 0.05; all z > 1.96 controlled for multiple comparisons with Benjamini-Hochberg procedure; n = 426). In line with our prior work these results show that the saccade-fixation cycle dynamically influences network interactions (*26*). These results show also that the amygdala-hippocampal connectivity is sensitive to other than emotional forms of saliency. They support the idea that this circuitry contributes to a more general process of evaluating biological significance of visual stimuli and enhances sensory processing through pathways to the visual cortex (*62–64*).

## Discussion

We developed closed-loop eye tracking gaze contingent stimulus presentation and combined it with iEEG recordings in surgical epilepsy patients to investigate the physiology of active vision. By systematically examining neural activity at different phases of the saccade-fixation cycle during active vision we made several new observations. First, we found that sensory inputs presented during active vision (i.e., following saccades) elicit stronger and faster neural responses. Active-passive differences include increased phase coherence in alpha/theta range and decreased power in the same range during active vision. Second, we observed that hippocampal neural excitability undergoes cyclical excitability fluctuations (aka oscillations) at a theta-alpha rate tied to the saccade-fixation cycle. Third, we found that different types of information in a sensory input are preferentially encoded at distinct phases across the saccade-fixation cycle. Fourth, we observed that active vision decreases alpha– and increase theta-range phase synchronization in the amygdala-hippocampal circuit. Overall, our results support a model in which the saccade-fixation cycle is a basic unit organizing sampling, coding and network transfer of visual information. These results highlight the importance of studying vision in the context of natural active sensing behavior. Several of these findings merit further emphasis.

### The saccade-fixation cycle is a basic unit organizing sampling of sensory information

Previous studies have demonstrated that saccades reorganize neural activity across a large network of areas spanning the visual hierarchy (*5*, *13–21*) up to the hippocampus (*24*, *25*, *28–31*, *65*) and extend into nonvisual sensory networks (*26*, *27*, *66*) and thalamic nuclei (*32*). Our recent work has demonstrated that saccades modulate neural excitability in the auditory and somatosensory systems, and that the intensity and direction of interactions between the Frontal Eye Fields and sensory areas change across the saccade-fixation cycle (*26*). The primarily top-down direction of interaction towards the end of fixation and during saccade is replaced by a bottom-up direction at the onset of fixation (*26*); this same type of alternation was also observed at the laminar circuit level in V1 (*23*). These studies suggest that the saccade-fixation cycle plays a role in organizing the uptake and global distribution of sensory information modulating large scale network interactions across the brain.

Here, we show that during active vision saccades elicit fluctuations to neural excitability at theta and alpha rates which organize encoding of sensory information across the saccade-fixation cycle. Our observation that excitability fluctuates systematically across the saccade-fixation cycle accords with a previous study in nonhuman primates (*23*), albeit with differences in the time course of excitability fluctuation between the studies. Here we show that hippocampal excitability increases from fixation onset throughout the time course of a typical fixation duration (∼200 ms). In contrast, V1 excitability decreased with time from the period immediately after fixation. We posit that these differences to excitability time course may arise due to saccade-related phase reset differently modulating neural excitability across the visual hierarchy. For example, lower level visual areas might be shifted to a high excitability state right after fixation while higher level areas, including the hippocampus, are shifted to a high excitability with a certain lag (*26*, *32*). Additionally, differences between these studies may result from the way excitability time course is estimated. Here, we used BHA to track excitability, while multiunit activity (MUA) was used by Barczak et al. (*23*). Our previous work has shown that BHA and MUA have different laminar generators and that BHA may be in part subthreshold to neural firing (*41*). This increased sensitivity of the BHA signal may account for the additional alpha modulations observed in our study. Overall, our study highlights the importance of examining neural dynamics across the saccade-fixation cycle. Including natural saccadic sampling conditions is important in understanding the fundamental organization of sensory processing and information coding. Critically, passive approximations to natural vision lack these dynamics that arise during active vision, potentially resulting in a distorted understanding of natural vision, particularly the interplay between top-down and bottom-up processes.

### Coarse-to-fine coding of information across the saccade-fixation cycle

We observed that sensory information is phase-coded during active vision, with visual qualities such as intensity and edges preferentially encoded at distinct phases of the saccade-fixation cycle. Specifically, field potentials closer to the saccade (i.e., 50 ms following fixation onset) contained more information about stimulus intensity. This was followed by information about edges, which was preferentially represented later in the cycle (i.e., 100 ms following fixation onset). This coarse-to-fine encoding of visual information aligns well with previous psychophysics findings that increased sensitivity to low spatial frequency (i.e., intensity) is strongest shortly after a saccade, followed by heightened sensitivity to high spatial frequencies (i.e., edges) later in the fixation period (*51–54*). Our results, in conjunction with these psychophysics (*51–54*) findings, suggest that during active vision, predominantly spatial activation of retinal input is transformed into a temporal code with distinct visual features preferentially encoded along the saccade-fixation cycle. Coarser elements (i.e., intensity) are represented closer to fixation onset, followed by finer elements (i.e., edges).

What mechanisms underlie this coarse-to-fine coding of visual information? One possibility is that both saccades and fixational eye movements (drifts and tremors) contribute to encoding different spatial information at each saccade-fixation cycle. At fixation onset, neurons are primarily stimulated by low-frequency changes driven by saccades, followed by high-frequency changes driven by fixational eye movements. The alternation between saccades and fixational drifts during active sensing cyclically exposes neurons to these different modulations (*1*, *12*, *51–54*). Supporting this interpretation, fixational eye movements have been found to attenuate low spatial frequencies and accentuate high frequencies (*51–54*). Alternatively, signals about intensity and edges may reach the hippocampus via pathways with different conduction times. For instance, edge representation may primarily involve slower parvocellular pathway projections through the ventral visual stream via the perirhinal and lateral entorhinal cortex to the hippocampus. Conversely, intensity representation may involve faster magnocellular pathway projections to the dorsal stream via the parahippocampal gyrus and medial entorhinal cortex to the hippocampus (*49*, *50*, *67*, *68*).

A recent study used dynamic machine learning algorithms to investigate the impact of low-resolution sensor movements, mimicking fixational drifts, on classifier performance within a static scene context (*69*). The authors found that incorporating dynamic sampling-like behavior into a convolutional neural network with recurrent connectivity significantly enhanced classifier performance compared to the initial performance. This computational study shows the importance of small fixational eye movements in encoding sensory information. Our results suggest that systematically varying the size of these displacements could further enhance encoding and classification performance. Implementing dynamic sampling-like movements with steps of different sizes (analogous to large saccades followed by small fixational eye movements) may improve classification performance by supporting the encoding of different types of information.

Altogether, previous psychophysical, computational, and our neurophysiological results support the notion that different types of information are encoded at various phases of the saccade-fixation cycle. Coarse elements of a visual scene are encoded closer to the saccade, while finer elements are encoded later in the fixation period. This phased encoding strategy highlights the intricate relationship between eye movements and visual information processing, offering new insights into the mechanisms of sensory encoding and their potential applications in machine learning and neural computation.

### Phase coding of information during active vision

While the phase coding observed across the saccade-fixation cycle in this study is a novel discovery, previous research has demonstrated that distinct phases of neuronal oscillations can carry different information about sensory inputs, including natural sounds (*70*, *71*), visual inputs (*72*, *73*), and memorized objects (*74*). Our findings indicate that neural activity in the hippocampus represent the spatial properties of visual inputs in a temporal code. It remains to be determined whether the hippocampus actively participates in this spatial-to-temporal transformation or whether this transformation occurs at earlier stages of sensory processing or results from the sampling behavior as discussed above. Our results support a model in which saccades reset the phase of ongoing neural fluctuations across a large network organizing information coding and inter-areal interactions.

### Do we see saccadic suppression in the hippocampus?

Previous studies have documented a loss of sensitivity in the magnocellular visual pathway, which is suppressed during saccades (*75*). Psychophysical research has shown that this suppression begins approximately 50 ms before saccade onset, peaks at the moment of movement, and persists for about 50 ms after the saccade, followed by post-saccadic facilitation (*76–78*). Electrophysiological recordings from motion-sensitive areas in both the dorsal (*79*, *80*) and the ventral stream, as well as in the medial temporal lobe (*13*, *25*) and lateral geniculate nucleus (*15*), align well with the temporal profile of perceptual suppression. Recently, we identified a similar process in the auditory system, exhibiting comparable temporal dynamics (*26*). In the current study, we observed analogous perisaccadic suppression in the human hippocampus, with stimuli presented at the shortest lag (i.e., 50 ms following saccade offset) eliciting the weakest BHA response. Similarly, a number of neurons have been identified in the human hippocampus that exhibit a comparable time course of perisaccadic suppression(*25*). Saccade-locked analyses of BHA in the anterior nuclei of the thalamus—areas directly connected to the hippocampus—also revealed an initial BHA suppression followed by a rebound effect (*32*). Collectively, these findings suggest that perisaccadic neural activity is suppressed in the hippocampus and its connected regions, mirroring the well-documented phenomenon of saccadic suppression in the visual system.

### Active vision influences network activity in the amygdala-hippocampal circuit

Synchronization of neural oscillations is regarded by many as a crucial mechanism for inter-regional communication (*61*, *81–83*). Notably, low frequency phase synchronization within the amygdala-hippocampal circuit is associated with memory encoding of emotionally salient information (*59*, *60*, *84*, *85*). Alpha phase desynchronization and theta synchronization in this circuit supports successful mnemonic discrimination (*60*). Here, we observed that active vision promotes alpha phase desynchronization followed by theta synchronization in the amygdala-hippocampal circuit. Our results show that this pattern of alpha-theta phase synchronization is not limited to mnemonic discrimination of emotional events but extends to other contexts, possibly reflecting a more general process of marking inputs of increased behavioral relevance (*62–64*). Our results suggest also that passive vision, by increasing alpha and decreasing theta phase synchronization in the amygdala-hippocampus circuit, may promote formation of false memories (*60*). While this exact idea remains to be tested some support for it was recently been reported (*86*).

Our results suggest also that selective phase reset may control propagation of network activity and support routing of information through brain circuitry. We observed phase reset during active vision in the hippocampus but detected no evidence of a similar effect in the amygdala. This selective phase reset transiently desynchronizes alpha-range activity in the circuitry. While the suggestion of gating by selective phase reset is novel a similar role has been attributed to local alpha power. For example, Jensen and Mazaheri (*87*) proposed that increasing alpha power in an area suppresses neural activity in that area but also disconnects that area with other regions effectively re-routing propagation of network activity (*88*). Our results suggest that phase may play a similar role with selective phase reset transiently disconnecting an area from a network.

### Potential limitations of our results

Could our results be explained by cognitive factors that are not specifically related to saccades? The study was designed to keep overall memory and attention loads comparable in both active and passive condition. The memory load was minimal. Both in active and passive condition participants needed to maintain no information other than the initial stimulus identity. Because repeated fixations to the same location were allowed, participants did not need to memorize locations previously fixated. Similarly, attention load was comparable across conditions. Participants needed to monitor presented stimuli and respond to infrequent probes (every 4-14 sampled uniformly; see Methods). We observed that overall accuracy was high and did not differ across active and passive condition though reaction times were shorter in active compared to passive condition. It is possible that the need to plan and execute a movement imposes a small additional cognitive load in active condition. However, an increased load in active condition would rather lead to prolonged reaction times and delayed ERPs onsets which is opposite to what we observed.

It is also unlikely that our results are explained by differences in visual stimulation across active and passive conditions. First, we used the same stimuli in both conditions. It is also unlikely that at longest intervals participants relocated their gaze before the image was displayed. The longest interval of 200 ms was selected to be below the range of typical saccadic latencies (*89*). Importantly, if saccades away from stimulus in the active condition were to explain the current results, one would expect this condition to elicits smaller neural responses compared to the passive condition, which is opposite to what we observed. Furthermore, such an interpretation does not account for changes to neural responses at shorter lags including differences between passive condition and shortest active intervals (see Fig. 3).

One potential limitation of the current study is that in the active condition consecutive stimuli are presented at different locations while in the passive condition consecutive stimuli are presented at the same location. While this cannot explain most of our results it is worth discussing this as a possible limitation of the study. First, note that overall gaze directions are balanced (i.e., the number of stimuli presented in all four gaze directions are comparable in both active and passive condition). Similarly, the retinotopic locations are comparable because in both active and passive conditions stimuli were presented foveally. It is the spatial location of consecutive stimuli relative to external cues (e.g., monitor frame, background behind the monitor, etc.) that differs between active and passive conditions. These differences cannot explain our observations about fluctuating neural excitability and information coding across the saccade-fixation cycle (Fig. 3 and 4) because these results come from active vision conditions entailing different time lags between fixation onset and stimulus presentation. In these conditions consecutive stimuli are always presented at different locations. Finally, one may imagine that two consecutive stimuli presented at different locations (even without the need to move eyes) elicit more phase aligned responses to the second stimulus, stronger BHA, weaker power and more aligned phase coherence int the amygdala-hippocampal circuit (Fig. 2 and 5). However, if these differences were solely explained by the spatial mismatch between the stimuli, it would be difficult to understand how phase synchrony (including higher phase synchrony preceding stimulus presentation) would survive the inter-stimulus-interval. Assuming saccades have no influence on neural response in active condition, one would need to assume that any synchrony survives inter-stimulus-interval jitter (composing of irregular fixation and saccade duration) which is very unlikely given the tight temporal alignment necessary for measuring phase synchronization. Altogether, we acknowledge that this possibility worth discussing, but we think that the influence of active vision is a more parsimonious explanation.

## Conclusion

Our findings demonstrate that saccades during active vision play a pivotal role in reorganizing neural excitability, input coding and network connectivity, thereby facilitating the processing of information. The saccade-fixation cycle thus represents a fundamental unit that orchestrates the sampling of visual information and its routing across neural networks. These results highlight the importance of investigating vision within the framework of natural active sensing behaviors.

## Methods

### Patients

Continuous eye tracking and iEEG data were recorded from 7 patients (mean age 29.4; age range 20 – 40; 4 female) implanted with electrodes for surgical treatment of refractory epilepsy. All recordings were performed at the Columbia University Medical Center in New York. The study was approved by the institutional review board at the Columbia University and all patients gave written informed consent.

### Task

We developed a closed-loop eye tracking task which allows to present an image in a gaze contingent manner with a total system delay below 10 ms. Participants performed active and passive visual search. In the active search condition, participants were initially presented with a small image (∼2.5 dva in each direction from the screen center) er) at the screen’s center. They were instructed to fixate on this image and memorize its details. After 1 second of continuous fixation, the image disappeared, prompting participants to move their eyes across four different pre-designated locations on the screen in search of images. Upon fixating on one of these locations, an image was promptly presented to their fovea for 17 ms duration. The screen remained uniformly gray during the visual search, ensuring that images were presented only after a saccade, with no indication of the expected fixation locations. Participants familiarized themselves with these locations during a training session prior to the experiment. At random intervals during the visual search, participants were prompted to indicate whether the last image they saw matched the initial image in the trial. Overall accuracy was consistently high and did not differ between active and passive conditions.

Each participant completed a total of 40 active search trials, with the number of images per trial varying between 4 and 14 (distributed uniformly). In half of the trials, the last image was repeated. Additionally, we systematically manipulated the fixation-stimulus interval by presenting images with one of four different time delays (50, 100, 150, or 200 ms) after the eye was detected in one of the designated locations. This manipulation aimed to assess cortical excitability at different time points following fixation onset.

In the passive search condition, the initial image appeared in one of the screen quadrants, mirroring the spatial locations used in the active condition. Participants were instructed to fixate on this image for 1 second, after which a sequence of images flashed in that location. Similar to the active condition, participants were prompted at random intervals to indicate whether the last presented image matched the first image in the trial. Participants completed a total of 40 passive search trials, with the number of images per trial ranging between 4 and 14 (matching the active search but in reverse order). As in the active condition, half of the trials included repeated images.

Active and passive trials were randomly interspersed throughout the experiment, with careful matching of image content and inter-stimulus intervals between conditions. Specifically, the inter-stimulus interval in the passive condition was derived from those in the active condition. Identical images were presented in both conditions, albeit in reverse order to prevent habituation.

Stimuli comprised faces (82) and places (83), with image-locked field potentials analyzed in both passive and active conditions for the purposes of this manuscript.

### Electrophysiology data acquisition and preprocessing

Intracranial EEG (iEEG) data were filtered between 0.3 and 500 Hz (Butterworth 4th order) and digitized at a sampling rate of 2 kHz with 16-bit precision using equipment from Blackrock Microsystems LLC, focusing on recordings from the hippocampi and amygdalae of epilepsy patients. Subsequently, the data underwent downsampling to 500 Hz for further processing. To enhance spatial specificity and minimize common noise and contributions from distant sources via volume conduction, we implemented a bipolar montage technique. This involved subtracting signals from neighboring contacts on each electrode shaft. Line noise was removed using band-stop filters (Butterworth 4th order) at 60 Hz, 120 Hz, and 180 Hz.

Spectral phase and power were then estimated on the continuous data using 3-cycle Morlet wavelets across frequencies ranging from 1 to 30 Hz in 1 Hz steps. The resulting complex-valued time series was rectified and squared to extract power. For the calculation of Broadband High-frequency Activity (BHA), frequencies spanning from 70 to 150 Hz in 2 Hz increments were analyzed. A sliding Hanning tapered window (150 ms) with 6 Hz spectral smoothing was employed, and the complex-valued signal was rectified and squared to derive power. The frequency dimension was subsequently averaged to generate a single vector representing BHA fluctuations.

The data were then segmented into three categories: (1) raw field potentials without any additional filtering for calculation of ERPs (following previous studies ERPs were rectified before averaging), (2) time-frequency complex-valued time series spanning 1-30 Hz, and (3) BHA time-series relative to stimulus presentation, with a temporal window of 1200 ms before and after stimulus onset. Similarly to our previous studies (*26*, *32*), epochs containing artifacts were defined as those in which the temporal difference of field potential exceeded 5 standard deviation of the trial mean.

Note that Fig. 2A (passive condition) and Fig. 3A (passive condition) present the same BHA data (n = 76), smoothed in two different ways. In Fig. 2 we used moving average with 50 samples kernel, while in Fig. 3 we used Gaussian smoothing kernel with standard deviation of 20 samples. These filters were only used for illustration purposes. All statistical analyses were performed on unfiltered data.

<u>Inter-trial phase coherence (ITC)</u> was quantified at each time-frequency point. We used single trial (i.e., stimulus locked) time-frequency complex-valued signal and previously described formula (*55*) with the following Matlab implementation:

ITC = abs(mean(exp(1i * angle(complex-valued signal))

<u>Generation of feature maps for RSA</u>. In our RSA analyses, we incorporated five distinct image qualities as defined by the Itti and Koch model (84, 85), namely edges, saliency master map, orientation, intensity, and color feature maps. To extract edges from the images, we initially converted each image from RGB format to 2-D grayscale. Subsequently, we employed Matlab’s Sobel algorithm implementation to detect edges in each image. Remaining features were extracted using the Graph-Based Visual Saliency (GBVS) Toolbox implementation of the Itti and Koch model (*85*). Specifically, applying the GBVS algorithm to our image dataset, we employed default parameters, including a saliency master map size of 19×19 pixels and feature channels encompassing orientation, intensity, and color. The GBVS algorithm extracts low-level feature maps (orientation, intensity, and color) from images in RGB format using biologically inspired filters. These feature maps are then integrated to compute a unified global saliency master map. Subsequently, stimulus distance matrices were generated for each feature by computing Euclidean distances between pairs of images. This process was performed individually for each feature, resulting in five distinct stimulus distance matrices, each corresponding to a different feature.

<u>Representational similarity analysis</u>. To test the hypothesis that varying amounts of information regarding visual features are encoded across different phases of the saccade-fixation cycle, we conducted a representational similarity analysis (RSA). This analysis aimed to assess the degree to which evoked patterns reflect low-level features within the stimulus space. To investigate the possibility of phase coding throughout the saccade-fixation cycle, we conducted separate analyses for stimuli presented at four distinct time intervals (i.e., 50, 100, 150, 200 ms relative to fixation onset). For each image, we extracted five distinct feature maps representing edges, saliency, orientation, intensity, and color. Subsequently, we computed Euclidean distances between pairs of stimuli for each feature, generating stimulus distance matrices for each feature and time lag. Similarly, we calculated the distance matrix of the neural data, representing pairs of evoked responses, using the same approach. To assess the extent to which neural data track visual features, we correlated (Spearman’s rank correlation) stimulus distance matrices with neural distance matrices for each feature and time lag separately (see Fig. 4A for the analysis overview).

If field potentials contained varying amounts of information about visual features across different phases of the saccade-fixation cycle, we anticipated non-uniform RSA results across the time lags. Furthermore, we hypothesized that distinct visual features might be preferentially encoded at specific phases of the cycle. Alternatively, field potentials might exhibit similar types and amounts of information across the cycle.

We applied RSA to single-trial electrophysiological data patterns, comprising concatenated Broadband High-frequency Activity (BHA) and Event-Related Potentials (ERPs) within a predefined time window (300-500 ms after stimulus presentation), corresponding to the period of strongest sensory elicited potential in the hippocampus (43, 44). To test for any differences in the degree to which the neural data reflects features in the stimuli space we used Kruskall-Wallis’ test comparing Fisher’s transformed correlation coefficients across the four lags. Following previous studies, we also tested whether Spearman’s correlation coefficients for each feature and each lag differed from zero (50, 51). We used a one-tailed Wilcoxon sign rank test because negative RSA are difficult to interpret and are often considered noise (50, 51). Note that we also reproduced all our effects using a two-tailed Wilcoxon sign rank test.

<u>Decoding analyses</u>. To further explore the dynamics of information processing across the saccade-fixation cycle, we employed a decoding approach. Our aim was to investigate how the amount of information about sensory input (i.e., stimulus category: faces vs. places) varies throughout the saccade-fixation cycle. Particularly, we focused on the hypothesis that the amount of information is not uniform across the cycle with different amounts of information regarding stimulus category encoded preferentially at some parts. We utilized the area under the curve (AUC) as a metric to quantify the amount of information extractable by a classifier from the data. To assess information encoding at various phases of the saccade-fixation cycle, we compared classifier performance across four distinct intervals relative to fixation onset (i.e., 50, 100, 150, or 200 ms following eye detection at designated locations).

Our rationale posited that if the amount of information varied across the cycle, classifiers trained and tested on data from specific phases would demonstrate non-uniform accuracy in decoding stimulus categories. Specifically, we aimed to predict high-level visual information (stimulus category: face vs. place) from hippocampal field potentials using a 7-fold cross-validation approach, coupled with PCA feature reduction. Data were centered around the peak hippocampal ERP response, occurring within the 300-500 ms post-stimulus time frame. To ensure the robustness of our findings, we employed three distinct binary classifiers: Gaussian Naive Bayes, linear discriminant analysis, and support vector machines with a linear kernel and a-priori parameter c set to default c = 1. This multi-algorithmic approach minimized dependence on any specific algorithm, enhancing the reliability and generalization of our results

<u>Connectivity analysis</u>. To quantify connectivity, we employed the Phase Locking Value (PLV). Similar to the ITC described above, we utilized single-trial, stimulus-locked complex-valued time-frequency signals and applied the previously described formula (55) with the following Matlab implementation:

PLV = abs(mean(exp(1i * (phase_x – phase_y))

Where phase_x and phase_y refer to the two complex-valued signals from two channels.

<u>Could the decrease in power explain the decrease in PLV?</u> There are two reasons why the current results on phase synchronization while, similarly to power, are influenced by our experimental manipulation, do not simply reflect changes in signal-to-noise ratio. First, fluctuations in signal-to-noise ratio would have instantaneous influence on the PLV values. In turn, we observe that power and PLV increase in passive condition both show a different time course and frequency composition. We observed strongest power increase in *<u>passive</u>*condition ∼ 400-600 in a relatively wide range of frequencies from 5 to 20 Hz. The PLV around that time was limited to lowest frequencies (∼1 Hz) and showed increase in active condition. Second, except for some extreme cases when power is very low, PLV and power appear mostly independent. For example, Muthukumaraswamy and Singh, 2011 observed that power spuriously influences PLV only at relatively low SNRs but the effects are negligible at higher signal-to-noise-ratios (SNRs). Similarly, another simulation was performed using empirical data to estimate the influence of power on PLV (Cohen, 2014: p. 341-342). Empirical neural time series from one or both electrodes were multiplied with a constant number akin to that performed by Muthukumaraswamy and Singh, 2011 but with realistic (i.e., empirical) baseline power values. These multiplications had little influence on PLV estimation. Altogether, these simulations suggest that power changes may influence PLV but only at very low SNRs and are negligible at higher SNRs (including empirical power). It is also worth mentioning that this later simulation was performed using scalp recordings which typically show lower SNRs compared to the intracranial recordings used in the current study. This again suggests that the current PLV results are relatively immune to power fluctuations.

<u>Statistics.</u> We employed non-parametric Wilcoxon signed-rank tests to compare the magnitude of field potentials, BHA, spectral phase coherence, and power across frequencies ranging from 1 to 30 Hz during active and passive viewing of the same set of images. To address multiple comparisons, we applied False Discovery Rate correction with the Benjamini-Hochberg procedure (86) across time for field potentials and BHA, and across time-frequency points for PLV and power.

Additionally, we conducted control analyses using linear mixed-effects models implemented with the lme4 package in R. A random intercept model was formulated as follows: response ∼ condition + (1| patient-id / channel-id). In this model, participant identification number was treated as a random effect, with channel identification nested within it. Experimental condition served as a fixed effect, and the magnitude of the signal of interest was the dependent variable (see below).

To quantify differences among four active vision conditions (lag 50, 100, 150, 200 ms; data presented in Fig. 3) we first applied a series of Kruskal-Wallis tests (p < 0.01 controlled for multiple comparisons using Benjamini-Hochberg method) to identify time windows during which visual stimuli evoked BHA of varying magnitudes across the saccade-fixation cycle. This analysis revealed three distinct windows: early (170-250 ms after stimulus onset), middle (340-350 ms after stimulus onset), and late (900-1000 ms after stimulus onset). To further characterize the specific BHA patterns within these windows, we conducted pairwise Wilcoxon tests between the conditions (Fig. 3 B, C, D respectively).

### Eye tracking data acquisition and preprocessing

Horizontal and vertical eye positions for both eyes were sampled at 300 Hz using a Tobii TX300 eye tracker. Calibration of the eye tracker was performed by capturing gaze fixations from five known target points. Following the calibration sequence, the calibration output underwent inspection, focusing on two key features. Calibration points lacking data or displaying dispersed gaze points around the calibration point were identified for recalibration. Once successful calibration was achieved, the experiment was initiated by the operator.

During the active search condition, an image was presented only when the eyes were relocated to one of the four specified regions (see Figure 1). Conversely, in the passive search condition, images were presented only if continuous fixation was maintained. If no eye was detected in the specified region, the experiment paused until participants fixated in the correct location. This method ensured that stimulus presentation during active search was preceded by an eye movement, while in passive search, it was preceded by continuous fixation. Breaking sustained fixation (passive condition) or not relocating gaze (active condition) halted the experiment until a participant fulfilled the requirement for the condition.

## Control analyses

### Phase coding of information across the saccade-fixation cycle: decoding approach

To investigate the potential for information coding across the saccade-fixation cycle, we employed a decoding approach. Information content was quantified as the accuracy (i.e., area under the curve; AUC) of a binary classifier predicting high-level visual information (stimulus category: face vs. place) from hippocampal field potentials (using Gaussian Naive Bayes; 7-fold cross-validation, PCA feature reduction, data centered at the strongest hippocampal ERP response: 300-500 ms post-stimulus). By training and testing separate decoders on evoked potentials elicited at four phases of the saccade-fixation cycle, we observed small but reliable variations in the amount of information learned at different time points across the saccade-fixation cycle (AUC = 0.50, 0.53, 0.51, 0.51; H(3) = 8.1, p = 0.04; Kruskal-Wallis test; N = 76). Decoding accuracy peaked 100 ms after fixation onset, surpassing levels observed at 50 and 200 ms lags (both p < 0.05; both z > 1.96), but not during the 150 ms lag (p = 0.059; z = 1.88; N = 76; sign rank test). Accuracy at the 100 ms lag also exceeded that of passive viewing conditions (p = 0.008, z = 2.63; N = 76; sign rank test). No significant differences in pre-stimulus baseline values were detected across lags (H(3) = 4.65, p = 0.2; Kruskal-Wallis test; N = 76). Comparing decoding accuracy in the post-stimulus window to pre-stimulus baseline, we observed increased accuracy from baseline to the 100 and 150 ms post-stimulus intervals (both p < 0.03, z > 2.14; N = 76), with no difference between baseline and post-stimulus intervals of 50 and 200 ms (both p > 0.10; z < 1.64; N = 76). To ensure the robustness of our findings, we conducted control analyses using linear discriminant analysis (H(3) = 17.58, p < 0.001; Kruskal-Wallis test; N = 76) and linear support vector machines (H(3) = 9.56, p = 0.02; Kruskal-Wallis test; N = 76), yielding similar results.

Overall, our findings suggest that field potentials in the human hippocampus encode small but reliable amounts of information about low– and high-level visual features, and that different features are preferentially represented at distinct phases of the saccade-fixation cycle. Our results suggest that human hippocampal field potentials encode small but reliable information about low– and high-level visual features, with different features preferentially encoded at distinct phases of the saccade-fixation cycle. These findings support a model in which information about sensory input is systematically linked to the saccade-fixation cycle.

### Control analyses – linear mixed effects models

In the primary analyses presented in the main text, we employed non-parametric rank-based statistics (i.e., Wilcoxon sign-rank tests), treating each electrode contact anatomically localized in the hippocampus as an independent unit of observation.

Various measures were implemented to mitigate the impact of clustering across channels within individual participants, including bipolar re-referencing, baseline correction, and averaging across multiple trials, all of which aimed to increase the independence of signals recorded from individual contacts.

To further ensure the robustness of our main results across potential dependencies, we conducted supplementary control analyses using linear mixed-effects models with the lme4 package in R. We formulated a random intercept model as follows: response ∼ condition + (1|patient-id / channel-id). In this model, we treated the participant as a random effect, with the channel as a nested variable and the experimental condition as a fixed effect, while the magnitude of BHA, early-ERP, and late-ERP, ITC-theta (3-7 Hz), ITC-alpha (8-12 Hz), and power (3-12 Hz) served as dependent variables.

To avoid over-parameterization and ensure model stability, we excluded random slopes and opted for a simpler model. The model was fitted with the REML estimator, and the significance of the fixed effect was determined using Wald F-tests with Kenward-Roger approximation for degrees of freedom. We inspected Q-Q plots and residuals versus fitted values to explore assumptions of the test (i.e., normal distribution and comparable variance of residuals). While we observed no major violations of the assumptions, in some cases (e.g., early-ERP, power), we noted small departures from normality. In such cases, we performed additional linear mixed-effects model analyses after log-transforming the response values prior to model fitting to better conform to the model’s assumptions. In summary, we reproduced our primary results in these control analyses, reinforcing the validity and reliability of our findings.

Specifically, using the linear mixed-effects models, we observed that BHA was increased in the active compared to the passive condition (F(1,75) = 14.542, p < 0.005). Because our initial inspection of grand averages showed two windows where ERPs may have differed, we performed separate analyses of the early-ERP (100-200 ms after stimulus onset) and late-ERP (500-600 ms after stimulus onset; see Fig. 2B). We observed stronger early-ERP (F(1,75) = 7.966, p < 0.01) and attenuated late-ERP (F(1,75) = 10.409, p < 0.01) in the active search condition. Additionally, we observed increased ITC in the theta (F(1,75) = 26.075, p < 0.01) and alpha (F(1,75) = 21.812, p < 0.01) frequency range during the active search condition. Power in the same theta/alpha frequency range was decreased in the active search condition (F(1,75) = 15.812, p < 0.01).

Inspecting residuals versus fitted values plots, we noted small violations of homoscedasticity. To ensure these do not influence our conclusions, we performed another test to control for possible clustering. To this end, we used the signed rank test with the Datta-Satten method to control for clustered data (87). The null hypothesis being tested using the Datta-Satten method is that the distribution of the paired difference of a randomly selected pair from a randomly selected cluster is different from zero. The Datta-Satten method accounts for the cluster size by assigning equal weight to each cluster instead of each paired difference; for example, a paired difference from a larger cluster will be assigned a smaller weight than a paired difference from a smaller cluster (88). Briefly, these control analyses reproduced the main results from our initial approach and the LME reported above for both the ITC in the theta (T = 19.632, p < 0.01; 2000 permutations) and alpha (T = 13.384, p = 0.02; 2000 permutations) range and power (T = 9.9563, p = 0.01; 2000 permutations).

## Funding

ML and CES are supported by a Silvio O. Conte Center Grant P50 MH109429, and by R01 DC012947. ML is supported by Polish National Science Center Grant 2022/45/B/HS6/04097.

## Author contributions

ML, CES designed the study. ML and ES collected data. ML, EE performed analyses. ML and CES wrote the manuscript.

## Competing interests

The authors declare no competing interests.

## Data and materials availability

All data needed to evaluate the conclusions in the manuscript are present in the paper and supplement.

Declaration of interests: The authors declare no competing interests.

## Notes

### Competing Interest Statement

The authors have declared no competing interest.

